# Molecular Characterization of SARS-CoV-2 N Protein Interfaces: Implications for Oligomerization, RNA Binding, and Phase Separation

**DOI:** 10.64898/2026.03.06.710084

**Authors:** Sneha G Bairy, Thazhe Kootteri Prasad, Yogeshwar Saravana Kumar, B Ganavi, S Shwetha, S Saranya, B Sriram Prakash, Vignesh Sounderrajan, Krupakar Parthasarathy, Neelagandan Kamariah

**Affiliations:** Centre for Chemical Biology and Therapeutics, Institute for Stem Cell Science and Regenerative Medicine and National Centre for Biological Sciences–TIFR, GKVK – Post, Bellary Road, Bangalore 560 065, India; Centre for Drug Discovery and Development, Sathyabama Institute of Science and Technology, Chennai, 600119, India

**Keywords:** SARS-CoV-2, Nucleocapsid protein, RNA interaction, homo-oligomerization, phase separation, Ribonucleoprotein complex, Nuclear magnetic resonance

## Abstract

The SARS-CoV-2 nucleocapsid (N) protein is central to genomic RNA recognition, condensation, and packaging, yet the molecular organization of its multivalent N–N and N–RNA interaction network involved in this process remains unclear. Here, we define the oligomerization and RNA-binding interfaces of the C-terminal domain (CTD) and its flanking intrinsically disordered regions (IDRs), the leucine-rich helix (LH) and the C-terminal IDR (C-IDR), using size-exclusion chromatography (SEC), cross-linking, mutational studies and NMR spectroscopy. We identify discrete oligomerization interfaces within the CTD and C-IDR that drive higher-order assembly, and show, through liquid–liquid phase separation (LLPS) and electron microscopy (EM), that C-IDR residues are essential for RNA-induced condensate formation. Moreover, the mapping of RNA-binding residues highlights Arg277 as a conserved determinant of CTD–RNA recognition. Notably, the two IDRs exert opposing regulatory effects on RNA binding, with the C-IDR enhancing and the LH attenuating CTD-RNA interactions. Together, these findings reveal how cooperative interfaces between the CTD and its flanking IDRs orchestrate N-protein oligomerization and RNA condensate formation and highlight potential intervention sites for disrupting SARS-CoV-2 ribonucleoprotein assembly.

## INTRODUCTION

Coronavirus disease (COVID-19) is a highly contagious illness caused by the SARS-CoV-2 virus. According to the WHO dashboard, there have been 779,142,314 confirmed cases and 7,111,892 reported deaths worldwide to date. In just the 30 days leading up to 15^th^ February 2026, nearly 48,187 new cases were reported (https://data.who.int/dashboards/covid19/cases). Several emerging variants—such as XEC, KP.3.1.1, LF.7.2.1, MC.10.1, NP.1, LP.8.1, and XFG—are spreading rapidly across the globe, underscoring the virus’s continued evolution (https://www.who.int/activities/tracking-SARS-CoV-2-variant). These developments highlight the need to better understand the molecular mechanisms that govern viral replication, assembly and pathogenesis. The SARS-CoV-2 life cycle within host cells involves critical stages, including unpackaging and packaging of viral genomic RNA (gRNA) during entry and egress [1, 2]. The viral genome is a ∼30 kb positive-sense single-stranded RNA (+ssRNA) that encodes 16 nonstructural proteins (nsps), 4 structural proteins, and 6 accessory proteins [3–5]. Assembly of new virions begins with encapsidation of the +ssRNA genome by the N protein into discrete ribonucleoprotein (RNP) complexes. These RNPs then bud into the ER–Golgi intermediate compartment (ERGIC), where they assemble with the membrane-associated structural proteins—Membrane (M), Envelope (E) and Spike (S) [6–8]. Each virion contains about 38 RNPs, each estimated to comprise 10-12 N proteins wrapping around 800 nucleotides [9–11]. Furthermore, specific genomic hotspots called Translational Regulatory Sequences (TRS) appear to serve as high-affinity binding sites for N, initiating genome packaging [5, 12, 13]. Despite significant progress, how a single copy of the viral genome is selectively packaged into one virion remains poorly understood [1, 3, 14].

The N protein is a central structural component of SARS-CoV-2, playing an indispensable role in RNA packaging, genome protection, and viral assembly. It is also a frequent locus of adaptive mutations that enhance viral replication, transmission, and fitness [15–19]. Recent studies suggest that N also functions as an RNA chaperone, efficiently unfolding and annealing RNA [20, 21]. Structurally, the N protein is 419 amino acids long, highly conserved among coronaviruses, and abundantly expressed in infected cells [22, 23]. N is composed of two folded domains: an N-terminal RNA-binding domain (NTD) and a C-terminal dimerization domain (CTD), connected by an intrinsically disordered central linker region (LKR) that is composed of serine/arginine (SR)-rich region, the leucine-rich helix (LH), and also flanked by N- and C-terminal disordered regions, namely N-IDR and C-IDR (Fig. 1A) [19, 24]. The N protein typically exists as a stable dimer in solution, mediated by the CTD, while the NTD is primarily responsible for RNA recognition. However, RNA-binding activity is not confined to the NTD alone. Multiple additional RNA-binding sites have been identified across the CTD as well as within IDRs, including the N-IDR, SR, LH and C-IDR [25–28]. Initial studies indicated that the NTD binds specifically to TRS sequences, whereas the CTD binds RNA non-specifically [13, 29, 30]. Later studies showed that the CTD is capable of binding RNA, and that its affinity for TRS increases further when it is combined with the C-IDR [25, 31–35]. Importantly, the C-IDR also mediates interactions between N and M proteins, a step required for final virion assembly [4, 36, 37].

**Figure 1.**
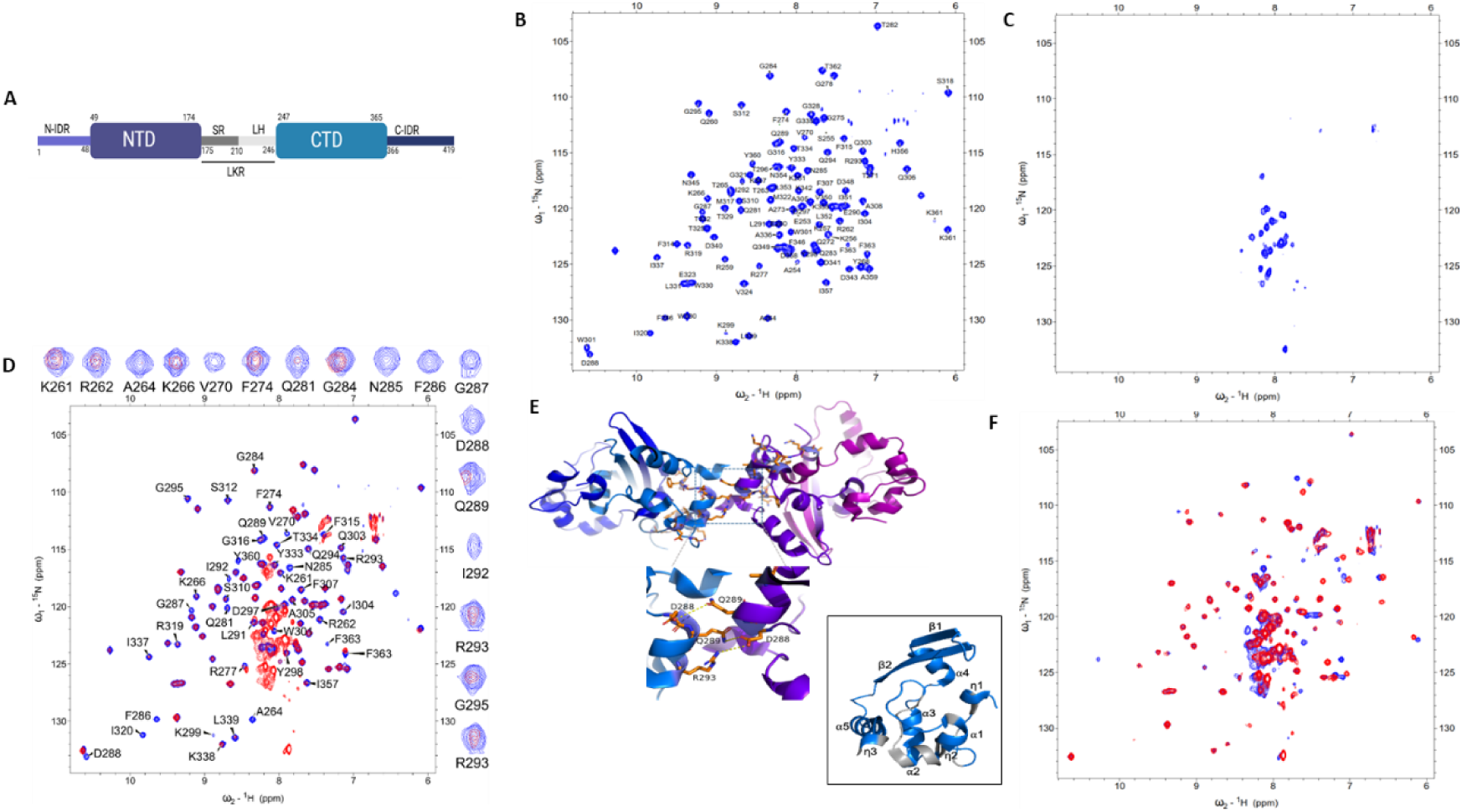
Structural organization of the N protein and identification of the CTD_247-365_ oligomerization interface in the presence of the C-IDR_366-419_. (**A**) Schematic representation of the SARS-CoV-2 nucleocapsid (N) protein domain architecture illustrating N-terminal intrinsically disordered region (N-IDR), N-terminal domain (NTD), the central linker region (LKR) containing Serine/arginine (SR) and Leucine-rich helix (LH) regions, C-terminal domain (CTD) and C-terminal intrinsically disordered region (C-IDR); HSQC NMR spectra of **(B)** ¹⁵N-labeled CTD_247-365_ ; (**C**) C-IDR_366-419_; (**D**) overlay of CTD_247-365_ (blue) and CTD+C-IDR_247-419_ (red). Residues in the CTD+C-IDR_247-419_ construct that exhibit significant peak intensity reduction relative to CTD_247-365_ are shown; (**E**) Residues within CTD_247-365_ mapped to the putative dimer–dimer interface, identified by reduced signal intensity associated with C-IDR induced oligomerization; (**F**) Overlay of HSQC spectra for CTD+C-IDR_247-419_ (blue) and CTD+C-IDR_DQR_ variant (red).

Besides RNA binding, homo-oligomerization is another key property of the N protein. Recent studies have highlighted the significance of an N protein fragment lacking residues 1–209 [N Δ(1–209)], encoded by the B.1.1 lineage that includes Alpha, Gamma, and Omicron variants [10, 38–40]. Despite the absence of the major RNA-binding NTD and N-IDR, this truncated form can still package RNA, form ribonucleoprotein (RNP) complexes, and assemble into virions [8, 26], indicating that critical binding interfaces are retained in the truncated form of N. Multiple studies emphasize the crucial role of LH region for the homo-oligomerization, liquid–liquid phase separation (LLPS), and viral assembly [10, 38, 41]. Moreover, oligomerization activity of the LH is regulated by phosphorylation of the adjacent SR region, with the unphosphorylated form favoring oligomerization and phosphorylation suppressing it [27, 42]. Despite these insights, the molecular details of N–N and N–RNA interactions mediated by the CTD and C-IDR remain unclear. Defining these interactions will be critical to fully delineate the N protein–RNA assembly pathway, its role in viral pathogenicity, and its potential as a therapeutic target.

In this study, we elucidate the molecular mechanisms underlying SARS-CoV-2 N protein homo-oligomerization and RNA binding, with a focus on the CTD and its associated IDRs, including the LH and the C-IDR. Using SEC, cross-linking, NMR spectroscopy, circular dichroism (CD) spectroscopy, mutational analysis, EM, and LLPS assays, we demonstrate that while the CTD and individual IDRs can each form stable dimers, their combination promotes tetramer formation and higher-order oligomers. We identify key residues in the CTD clustered around the α2-helix and the C-IDR residues covering 390 to 408, along with contributions from the LH region, as critical mediators of N-protein homo-oligomerization. Disruption of either the CTD or C-IDR interface perturbed LLPS, with the C-IDR truncation exerting a strong LLPS inhibition. Together, these findings establish that the CTD, C-IDR, and LH work together through distinct interfaces to drive N homo-oligomerization. Moreover, the NMR titrations coupled with mutational analysis further reveal RNA-binding sites in the CTD, with conserved residue R277 playing a central role. Both the C-IDR and LH bind RNA, yet they have contrasting effects on RNA binding to CTD. The C-IDR enhances CTD–RNA interactions, whereas the LH reduces CTD–RNA recognition. These findings suggest that IDRs play a regulatory role in shaping the RNA-binding specificity of the CTD. These key interfaces present promising targets for interfering with SARS-CoV-2 assembly.

## MATERIAL AND METHODS

### Oligonucleotides and nucleotides

Primers for cloning, single-stranded RNA derived from the viral genomic transcriptional regulatory sequence (TRS: 5′-AAACGAAC-3′), its DNA analog and the 32mer RNA from the 3’ S2M region of SARS-CoV-2 genome (5’-CGAGGCCACGCGGAGUACGAUCGAGGGUACAG-3′) [31] for biophysical experiments were purchased from Eurofins. RNA/DNA was reconstituted in RNAse-free water to 5 mM or 10 mM final concentration and stored at –80°C in small aliquots, which was thawed and used immediately. The full-length Nucleocapsid (N) gene of SARS-CoV-2, carrying a C-terminal 6×His tag, was obtained from Addgene (Plasmid ID: 162789).

### Construction of expression vectors

The N gene was cloned into the pET28a vector using a ligation-independent cloning method [43], introducing an N-terminal 6×His tag followed by a TEV protease cleavage site, while removing the original C-terminal His tag. Mutant and truncated N protein constructs were generated through site-directed mutagenesis and ligation-independent cloning, using primers listed in (Supplementary Table 1). All the constructs produced for the study are listed in (Supplementary Table 2). An N-terminal SUMO tag was added to the LH+CTD+C-IDR_210–419_ and LH+CTD_210–365_ constructs to enhance solubility. All recombinant plasmids were transformed into *E.coli* DH5α for amplification, and the coding sequences of all constructs were confirmed by Sanger sequencing.

### Protein expression and purification

The constructs were transformed into *E. coli* BL21(DE3) cells for protein expression. A single transformed colony was inoculated into 10 mL of LB medium containing kanamycin and grown overnight at 37°C with shaking at 200 rpm. This primary culture was then used to inoculate 1 L of LB medium, which was grown at 37°C until the optical density at 600 nm reached 0.8. Protein expression was induced with 0.25 mM IPTG, and the culture was incubated overnight at 18°C. Cells were harvested by centrifugation at 6000 rpm for 15 minutes at 4°C and resuspended in 40 mL of lysis buffer containing 25 mM Tris (pH 7.5), 5 mM MgCl₂, 1 M NaCl, 2% glycerol, 0.1 mM PMSF, and 5 mM β-mercaptoethanol. Cell lysis was performed on ice using a Qsonica sonicator (12 mm tip). The lysate was clarified by centrifugation at 17,000 rpm for 45 minutes at 4°C, and the supernatant was collected for purification.

The protein was first purified using Ni-NTA affinity chromatography. When the A₂₆₀/A₂₈₀ ratio exceeded 0.8, an additional heparin purification step was performed. For heparin chromatography, the protein sample was diluted threefold in 25 mM Tris (pH 7.5) buffer before loading onto the column, and eluted using a NaCl gradient (up to 1 M) in the same buffer. Fractions containing the target protein were pooled together and treated with TEV protease during overnight dialysis in size-exclusion chromatography (SEC) buffer, followed by a second Ni-NTA affinity step to remove TEV protease and uncleaved protein. The flow-through was concentrated and subjected to gel filtration using a Proteosec Dynamic 3–70 kDa 16/60 column both equilibrated with 25 mM Tris (pH 7.5), 300 mM NaCl, and 2% glycerol. For LH+CTD+C-IDR_210–419_ and LH+CTD_210–365_ constructs, the buffer contained 50mM Hepes (pH 7.5), 500 mM NaCl and 10% glycerol. The final peak fractions were concentrated to approximately 2 mg/mL and snap-frozen for storage.

### Molecular weight estimation using size exclusion chromatography

Protein molecular weight standards—ribonuclease (13.7 kDa), ovalbumin (43 kDa), conalbumin (75 kDa), and the BRCT domain of BRCA1 (27 kDa)—were used to calibrate the size-exclusion chromatography (SEC) The column void volume (V₀) was determined using Blue Dextran 2000. Each standard was dissolved in 500 µL of buffer (25 mM Tris, pH 7.5, 300 mM NaCl, 2% glycerol) and injected onto a Proteosec Dynamic 3–70 kDa 16/60 gel filtration column and for full length constructs γ-Globulin (150kDa) was added to the above standards and were used to calibrate the size-exclusion chromatography (SEC) Superdex S12 10/300 gel filtration column pre-equilibrated with the same buffer. The K_av_ value was calculated by the equation K_av_ = (V_e_-V_o_/V_c_-V_o_), where V_e_ is the elution volume, V_o_ is the void volume, and V_c_ is the column volume. K_av_ values were then plotted against the logarithm of the molecular weights of the standards, and the resulting calibration curve was used to determine the molecular weights of the constructs analyzed [44].

### Glutaraldehyde cross-linking

In a total reaction volume of 10 µL, protein at a concentration of 0.2 mg/mL was mixed with 0.3% glutaraldehyde (from a 5% stock) to initiate the crosslinking reaction, which was allowed to proceed at room temperature for 30 minutes. The reaction was quenched by adding Tris-HCl to a final concentration of 100 mM (from a 1 M stock), and the samples were subsequently analyzed by 12% or 15% SDS-PAGE.

### Nuclear magnetic resonance

For nuclear magnetic resonance (NMR) analysis, ^15^N-labeled proteins were produced in *E. coli* BL21(DE3) grown in M9 medium containing 0.5 g/L of ^15^NH_4_Cl. All ^15^N-labeled constructs were purified following the same protocol as the unlabeled proteins, with the NMR buffer containing 20 mM Tris-HCl (pH 7.5), 150 mM NaCl, and 1 mM DTT. For LH+CTD+C-IDR_210–419_ and LH+CTD_210–365_ constructs, proteins were eluted in the same buffer as the unlabeled proteins and subsequently buffer-exchanged into NMR buffer using a 10 kDa centrifugal filter. All NMR experiments were performed on a 600 MHz Bruker Avance III HD spectrometer equipped with a cryoprobe head, processed with Bruker TopSpin software, and analyzed with NMRFAM-SPARKY [45]. Typical ^1^H-^15^N TROSY-HSQC spectra were obtained at 298 K with 2048 points and 128 t1 increments, 48 scans per t1 point, and a 1.0 s recycle delay with a sweep width of 9615 Hz (^1^H) and 2128 Hz (^15^N). Backbone amide resonances were assigned based on previously reported structures of N_251–364_ (BMRB ID 50518 [46] and N_251–419_ (BMRB ID 51709 [35]. Titration experiments were performed using TRS RNA , its DNA analog and 32-mer stem loop RNA , by adding 5 mM or 10 mM stocks to the protein at molar ratios of 1:0.05, 1:0.5, 1:1, 1:2 and 1:5. Additionally, titration of CTD_247–365_ with the C-IDR_366–419_ was performed by adding unlabelled C-IDR_366–419_ in an equimolar ratio.

### CD spectroscopy

Circular dichroism (CD) spectra were acquired using a Jasco J-1500 CD spectrometer. Measurements were performed on 220 µl C-IDR_366–419_ samples at the indicated concentrations in buffer containing 50 mM Tris (pH 7.5), 150 mM NaCl, and 1 mM DTT, using a 1 mm pathlength quartz cuvette with a step size of 1 nm and averaging six accumulations. Secondary structural elements of C-IDR_366–419_ were estimated from CD spectra using BeStSel (https://bestsel.elte.hu/index.php) [47].

### Negative staining and transmission electron microscopy

A 10 µM FL-N and mutant proteins were mixed with an equimolar amount of an 8-mer TRS RNA in a final buffer of 12.5 mM Tris (pH 7.5), 75 mM NaCl, and 0.5% glycerol. A 2 µL aliquot of this mixture was applied to a glow-discharged, carbon-coated copper grid and incubated for 2 minutes to allow adsorption. Excess liquid was removed using Whatman filter paper, and the grid was briefly washed with 5 µL of water. Staining was performed with 2% (w/v) phosphor tungstic acid (PTA) for 10 seconds. After blotting, the grids were air-dried and imaged on a TALOS F200C G2 transmission electron microscope operating at 200 kV.

### Phase separation experiments using confocal microscopy

To assess phase separation in the absence of RNA, 30 µL of a 5 µM protein solution was directly dispensed into wells of a 96-well Eppendorf imaging plate. For RNA-induced phase separation, 5 µM protein was mixed with an equimolar concentration of TRS RNA or 32-mer RNA in a final buffer containing 12.5 mM Tris-HCl (pH 7.5), 75 mM NaCl, and 0.5% glycerol, yielding a total reaction volume of 30 µL. Samples were imaged at designated time points using an Olympus FV3000 confocal laser scanning microscope equipped with a 60× oil-immersion objective.

## RESULTS

### The IDR region flanking the CTD facilitates homo-oligomerization

The formation of large homo-oligomers by the N protein is essential for RNA packaging and viral assembly [12, 48]. The CTD, together with its flanking IDRs, is known to mediate homo-oligomerization [49]. However, elucidating its oligomeric behavior and the underlying molecular details is critical for understanding its role in RNA packaging. To investigate this, first, we systematically analyzed the oligomeric states of various CTD constructs incorporating flanking IDRs, including CTD_247–365,_ LH+CTD_210-365_, CTD+C-IDR_247–419_, LH+CTD+C-IDR_210-419_ alongside the full-length (FL) N protein N_1-419_. These constructs were expressed in *E. coli* and purified to homogeneity. SEC based molecular mass estimates revealed that FL-N_1-419_ and CTD_247-365_ primarily exist as dimers. In contrast, CTD_247-365_ incorporating additional IDR regions such as LH+CTD_210-365_, CTD+C-IDR_247-419_, and LH+CTD+C-IDR_210-419_ were capable of forming homo-tetramers (Supplementary Table 3). The crosslinking experiments indicated that CTD_247-365_ primarily forms dimers, accompanied by smaller populations of monomers, tetramers, and higher-order oligomers (Supplementary Fig. S1B). In contrast, LH+CTD_210-419_, CTD+C-IDR_247-419_, and LH+CTD+C-IDR_210-419_ formed notable amounts of dimers and tetramers. These constructs also produced substantial populations of heterogeneous higher-order oligomers. Notably, the homo-oligomerization pattern of CTD containing flanking IDRs is comparable to the oligomerisation profile of FL-N which similarly forms substantial amounts of dimer, tetramer and higher-order oligomers (Supplementary Fig. S1B). Together, these results indicate that the CTD, in combination with its flanking IDRs, strongly promotes homo-oligomerization.

### Identification of homo-oligomerization Interfaces in the CTD Domain

To further investigate the oligomerization interfaces of CTD and its flanking IDR regions, the ^15^N-HSQC spectrum of CTD_247–365,_ CTD+C-IDR_247-419_, C-IDR_366–419,_ LH+CTD_210-365_, LH+CTD+IDR_210-419_ were measured. The CTD_247–365_ spectra are well-dispersed, and resonance assignments were consistent with previously published data [46] (Fig. 1B). In contrast, C-IDR_366–419_, exhibited narrowly dispersed HSQC-spectra, characteristic of intrinsically disordered regions (Fig. 1C). Notably, the CTD+C-IDR_247–419_ spectra displayed characteristic features of both CTD and C-IDR regions. A detailed comparison of the HSQC spectra of CTD_247–365_ and CTD-IDR_247–419_ reveals that while peak positions of well-dispersed CTD peaks remain unchanged, the peak intensity of many of the CTD residues is significantly reduced in the CTD+C-IDR_247–419_ spectra (Fig. 1D). The observed line broadening and loss of CTD peaks are likely due to self-association/oligomerization of CTD+C-IDR_247 419_, as supported by SEC and cross-linking studies, also consistent with reports [31, 46, 50–52]. The reduction in peak intensity varies among residues, with a cluster—including K261, R262, A264, K266, V270, F272, F274, Q281, G284, N285, F286, G287, D288, Q289, I292, R293, G295, and D297—showing strong attenuation (Fig. 1D). These residues are localized in the N-terminal region of the CTD, mainly in α2 helix and its spanning η1, α1 and η2 helices and the loops connecting them (Fig. 1E). In addition, residues Y360 and F363 at the C-terminal end of the η3 helix are strongly affected, spacing closely with the α2 helix (Fig. 1E), implying a potential role of these regions in mediating homo-oligomerization.

Notably, the structural analysis of the dodecameric CTD_247–365_ [53] reveals a dimer–dimer interface involving residues whose intensity is strongly reduced in the CTD+C-IDR_247–419_ spectra. In this arrangement, the D288, Q289, and R293 residues in the α2 helix form a critical interaction network, with D288 from one dimer interacting with R293 and Q289 of an adjacent dimer. These findings suggest that residues in the CTD α2 helix, observed in crystal packing contacts, may form a viral assembly interface consistent with the classical helical model of coronavirus RNPs reported previously [7, 54]. To validate the involvement of these CTD residues in oligomerization, we generated and purified a triple mutant CTD+C-IDR_DQR_ in which D288, Q289, and R293 were substituted with alanine. The SEC analysis revealed that the CTD+C-IDR_DQR_ mutant eluted later than the CTD+C-IDR_247-419_, suggesting a weakened ability to form a stable homo-tetramer (Supplementary Table 4) (Supplementary Fig. S2A). However, a comparison of the CTD+C-IDR_DQR_ spectra shows no significant differences, as many CTD peaks remain missing in the mutant, similar to the wild-type CTD+C-IDR_247–419_ (Fig. 1F). This suggests that mutation of these residues in the CTD alone is insufficient to disrupt CTD+C-IDR homo-oligomerization.

### C-IDR region involved in the homo-oligomerization

SEC analysis of the purified C-IDR_366–419_ revealed that it predominantly forms a dimer, as reported (Supplementary Table 3) (Supplementary Fig. S2B) [49]. Cross-linking experiments further indicated that the C-IDR_366–419_ primarily exists as a monomer, with weak dimer formation with no significant levels of higher-order oligomers, suggesting the C-IDR alone has a weak propensity for homo-oligomerization (Supplementary Fig. S1B). To determine whether direct interactions between the C-IDR and the CTD contribute to stable tetramer formation, we conducted additional interaction studies. HSQC spectra of CTD_247–365_ upon titration with C-IDR_366–419_ revealed no detectable interaction (Fig. 2A).

**Figure 2.**
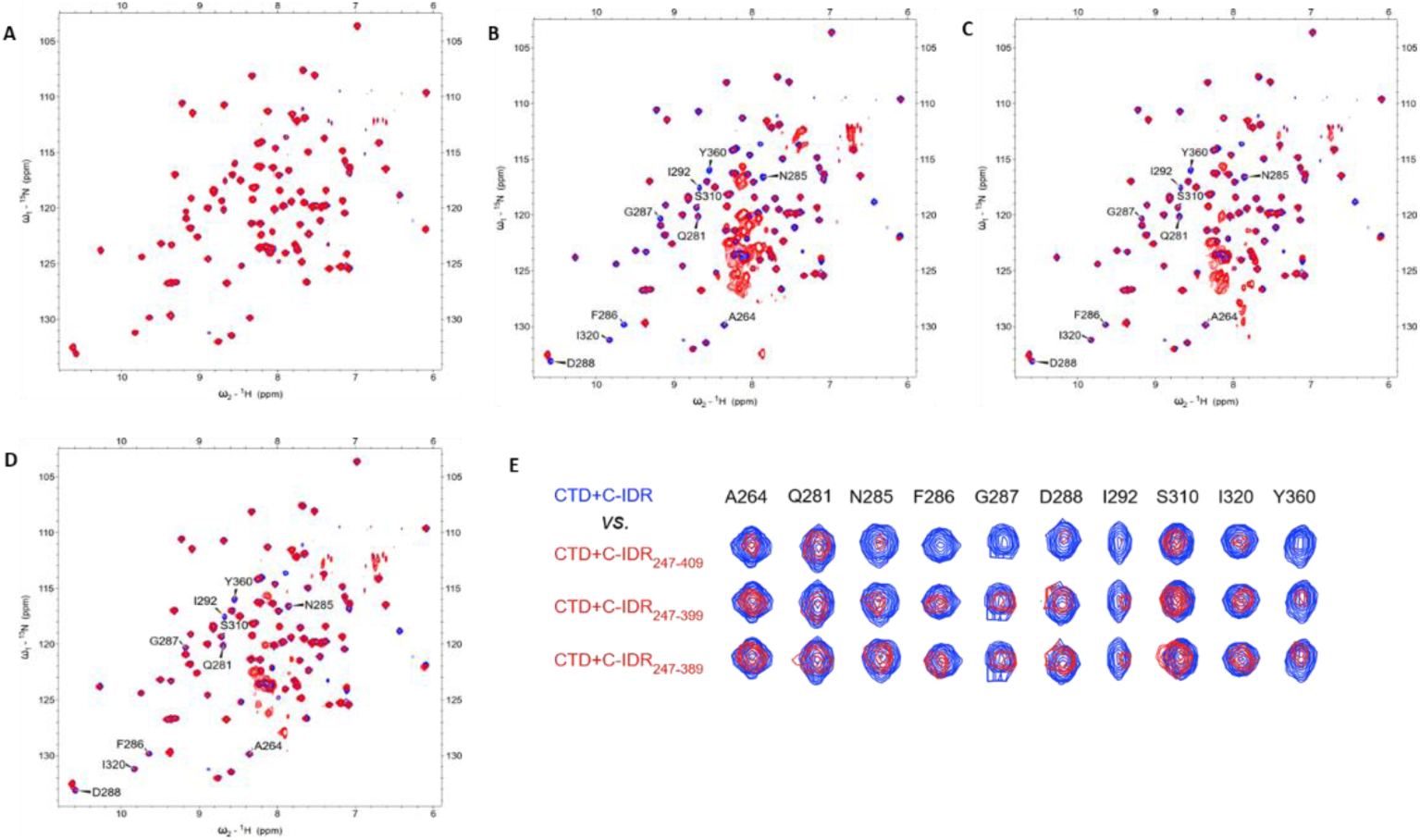
Mapping key C-IDR_366-419_ residues required for oligomerization. (**A**) HSQC NMR spectra of ¹⁵N-labeled CTD_247-365_ in the absence (blue) and presence (red) of unlabelled C-IDR_366-419_ at a 1:1 molar ratio; **(B-D)** Overlay of HSQC spectra of ¹⁵N-labelled CTD (blue) with , CTD+C-IDR_247-409_ (**B**, red), CTD+C-IDR_247-399_ (**C**, red) and CTD+C-IDR_247-389_ (**D**, red); (**E**) Comparison between CTD+C-IDR and CTD+C-IDR truncated constructs shows notable changes in signal intensity, indicating a progressive reduction in oligomerization as the C-IDR region is stepwise truncated.

To characterize the specific C-IDR segment involved in homo-oligomerization, we systematically deleted C-terminal residues in the CTD+C-IDR_247-419_ and purified variant deletion mutants such as CTD+C-IDR_247–409_, CTD+C-IDR_247–399_, CTD+C-IDR_247–389_ and CTD+C-IDR_247–379_. The SEC analysis showed that CTD+C-IDR_247–409_ eluted similarly to CTD+C-IDR_247–419_ forming a tetramer, whereas CTD+C-IDR_247–399_ eluted significantly later, indicating reduced oligomerization. Similarly, CTD+C-IDR_247–389_ and CTD+C-IDR_247–379_ also showed reduced oligomeric propensity compared to CTD+C-IDR_247-419_ and CTD+C-IDR_247–409_ (Supplementary Table 4) (Supplementary Fig. S2C). Moreover, the HSQC spectrum of CTD+C-IDR_247–409_ closely resembles that of CTD+C-IDR_247-419_, showing reduced peak intensity of well-dispersed CTD residues that are part of the oligomerization (Fig. 2B, E). Notably, the well-dispersed CTD peaks begin to reappear in the spectra of CTD+C-IDR_247–399_ and CTD+C-IDR_247–389_ (Fig. 2C, D, E). This suggests that residues spanning residues 390–409 exert a more pronounced effect on CTD+C-IDR_247-419_ homo-oligomerization compared to the segment containing the C-terminal tail residues 409–419 and the CTD+C-IDR_DQR_ mutant. The presence of highly polar and charged residues in the interface (390 QTVTLLPAADLDDFSKQLQQ 409) suggests that electrostatic interactions play a key role in driving homo-oligomerization.

To assess the helicity, the CD spectra of purified C-IDR_366–419_ were measured with increasing concentrations. Spectra measured at 50 µM indicated that the construct is predominantly disordered, displaying a strong negative peak near 200 nm characteristic of a random coil (Supplementary Fig. S3). Increasing the protein concentration (100 and 200 µM) resulted in gradual changes in the spectral profile, with the appearance of a shallow minimum near ∼222 nm. At 400 µM, two clear minima were observed near ∼208 and ∼222 nm, indicative of an increased helicity. Quantitative estimation of secondary structural elements using BeStSel further supported this trend, confirming an increase in α-helical content at higher concentrations (Supplementary Table 5). Together, these observations suggest that self-interaction of C-IDR promotes a concentration-dependent increase in secondary structure formation.

### Homo-oligomerization Interface of the CTD Modulated by LH

The SEC analysis confirms the stable tetramer formation of the LH+CTD_210-365_ (Supplementary Table 3) (Supplementary Fig. S1A). Cross-linking studies showed that, in addition to dimers and tetramers, LH+CTD_210-365_ has a strong propensity to assemble into higher-order oligomers. The HSQC spectrum of LH+CTD_210-365_ displayed well-dispersed peaks that closely resemble those of the isolated CTD, with only a few additional peaks in the narrow spectral region, likely arising from the disordered LH region (Fig. 3A). Notably, while a few CTD peaks showed reduced intensity compared to the isolated CTD, this effect was less pronounced than the extensive peak loss observed for CTD in the CTD+C-IDR_247-419_ spectra. The affected residues, such as K257, V270, T271, R277, Q281, F286, G287, I292, and Y360, cluster primarily within the α1 and α2 helices and the connecting loop of the CTD structure (Fig. 3B).

**Figure 3.**
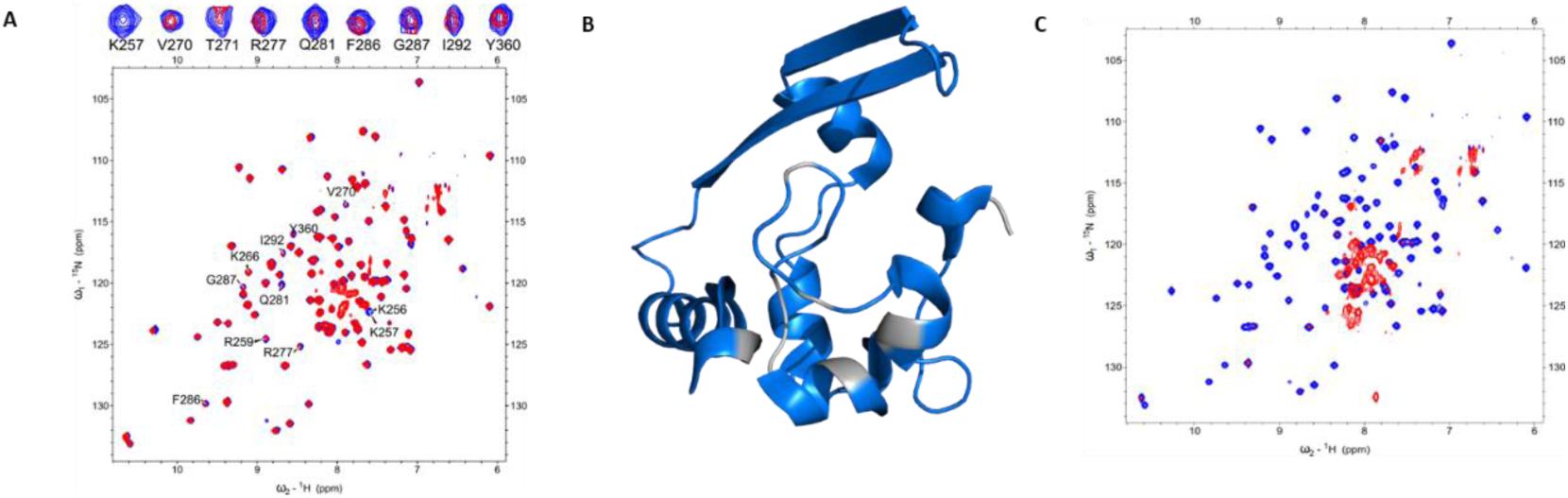
Identification of the CTD_247-365_ oligomerization interface in the presence of the LH. (**A**) HSQC spectra of ¹⁵N-labeled CTD_247-365_ (blue) and LH+CTD_210-365_ (red). Residues showing differences in peak intensity between CTD_247-365_ and LH+CTD_210-365_ are highlighted; (**B**) Mapping of CTD residues exhibiting reduced signal intensity in the LH+CTD_210-365_; (**C**) Overlay of HSQC spectra for CTD_247-365_ (blue) and LH+CTD+C-IDR_210-419_ (red), indicating a high degree of oligomerization in the LH+CTD+C-IDR_210-419_, evidenced by near-complete loss of detectable peaks.

Previous studies have shown that the LH domain can dimerize and form tetramers in the presence of the CTD [27]. The pronounced loss of LH peaks in spectra indicates that the LH domain plays a dominant role in driving homo-oligomerization, consistent with earlier findings that hydrophobic residues within the LH helix (residues 219–230) are central to the homo-oligomerization process [55, 56]. Our results further suggest that the LH domain serves as the primary driver, with the CTD interface playing a comparatively less extensive role in the oligomerization of LH+CTD_210-365._

By contrast, the HSQC spectrum of LH+CTD+C-IDR_210-419_ showed a complete loss of well-dispersed CTD peaks, with signals restricted to the narrow region corresponding to the disordered IDRs (Fig. 3C). This finding, together with cross-linking data (Supplementary Fig. S1B), indicates that LH+CTD+C-IDR_210-419_ undergoes an even higher degree of homo-oligomerization.

### Mapping the RNA-binding interface of the CTD

The N-protein contains multiple RNA-binding sites distributed across its two folded domains, NTD and CTD, as well as the flanking IDRs [25–27]. While RNA binding by the NTD has been well characterized, studies investigating the RNA-binding interface of the CTD remain limited, and its role in RNA packaging is not fully understood. To probe the RNA-binding interface of the CTD and assess the influence of flanking IDRs, we recorded HSQC spectra of CTD_247–365_, CTD+C-IDR_247–419_, and LH+CTD_2-10–365_ in the presence of an 8-mer single-stranded RNA (ssRNA) that contains the conserved transcription regulatory sequence TRS core (5’-ACGAAC-3’), flanked by two additional adenosines at the 5’ end [25]. The 8-mer TRS has previously been shown to bind CTD_247–365_ and CTD+C-IDR_247–419_ [25](Fig. 4A,B). We titrated CTD_247–365_ with a 32-mer stem-loop RNA. Even at a low protein:RNA molar ratio of 1:0.05, many resonances disappeared, consistent with the formation of higher-order oligomers (Supplementary Fig. S4). Severe line broadening and signal loss prevented residue-specific analysis with the stem-loop RNA.

**Figure 4.**
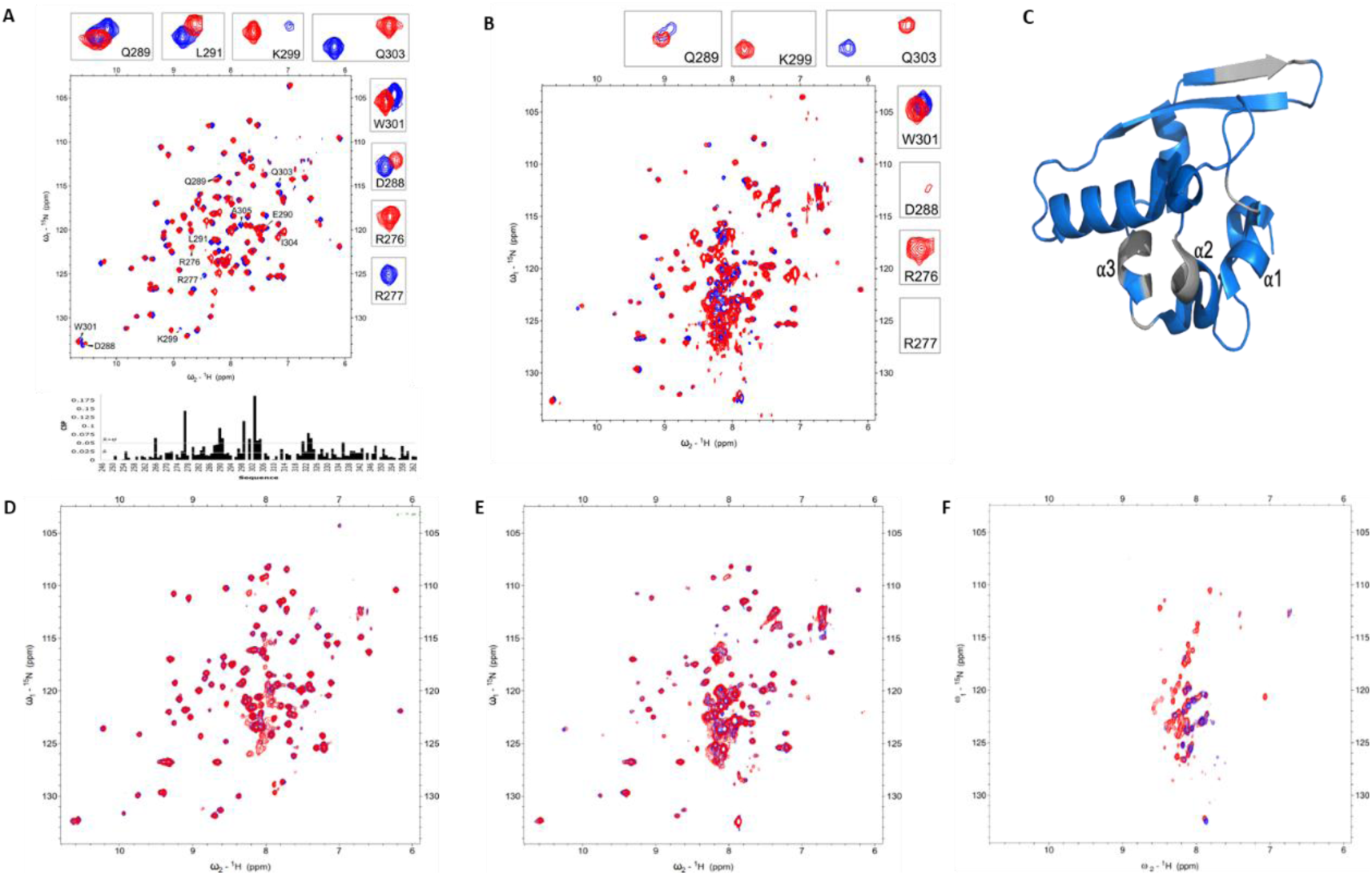
RNA binding interface of CTD in the presence of TRS RNA. HSQC NMR spectra of ¹⁵N-labeled constructs recorded in the absence (blue) and presence (red) of TRS RNA at the indicated protein:RNA molar ratios: (**A**) CTD_247-365_ (1:2); (**B**) CTD+C-IDR_247-419_ (1:1); (**C**) Structural mapping of RNA binding interface on the CTD; (**D**) CTD_R277A_ (1:2); (**E**) CTD+C-IDR_R277A_ (1:2); (**F**) C-IDR_366-419_ (1:1).

NMR titration experiments of CTD_247–365_ with 8-mer TRS RNA revealed significant chemical shift perturbations (CSPs), predominantly in the N-terminal region spanning residues 270–310 of the CTD (Fig. 4A). Within this segment, residues R277, R276, D288, Q289, E290, L291, K299, W301, Q303, I304, and A305 form a distinct cluster in the CTD structure (Fig. 4C). Among these, R277 was particularly notable: its peak, clearly visible in the apo spectrum, disappeared upon RNA binding. Even at lower equimolar TRS concentration, R277 exhibited a pronounced perturbation, underscoring the critical role of this positively charged residue in RNA recognition. Notably, R227 is highly conserved across related betacoronaviruses (Supplementary Fig. S5). In contrast, the R276 peak—absent in the apo form—emerged upon RNA addition, indicating conformational or dynamic rearrangements linked to binding. The other perturbed residues (E290, D288, Q289, L291, K299, W301, Q303, I304, and A305) localize mainly to the α2 and α3 helices, together with R277 and R276 in the α1–α2 connecting loop, forming the core of the RNA-binding interface (Fig4C). A secondary, more distal region showing weaker perturbations included residues G321–T325 (Fig. 4C). To further validate the RNA-binding interface as well as the critical interaction associated with the residue R277, we introduced an R277A mutation in both CTD_247–365_ and CTD+C-IDR_247–419_ constructs. RNA titration of the mutant proteins produced no measurable CSPs, confirming the identified binding interface and underscoring the critical role of R277 in RNA recognition (Fig. 4D,E). Whereas previous studies suggested RNA binding within the highly positively charged groove at the CTD dimer interface [53], our NMR titration results instead point to a moderately positively charged pocket, overlapping with the proposed homo-oligomerization interface, as the principal RNA-binding site (Supplementary Fig. S6). These findings further suggest that the CTD dimer may accommodate two RNA molecules, one bound to each monomer.

### The presence of the C-IDR enhances RNA binding of CTD and exhibits RNA specificity

Compared with CTD_247–365_, titration of CTD+C-IDR_247–419_ with 8-mer TRS ssRNA revealed a broadly similar interaction pattern (Fig. 4B). However, the CSPs for CTD+C-IDR_247–419_ at a 1:1 protein: RNA ratio were more pronounced than those observed for CTD_247–365_ even at a 1:2 ratio. In addition, equimolar TRS RNA induced the appearance of new peaks in the otherwise poorly dispersed spectrum of C-IDR_366–419_ (Fig. 4F), indicating its interaction with RNA. Due to spectral limitations, however, the specific RNA-contacting residues in this region could not be mapped. Nonetheless, the stronger RNA binding observed for CTD+C-IDR supports a cooperative contribution of both CTD and C-IDR to RNA recognition [25].

Previous studies have reported that CTD can interact non-specifically with both ssDNA and dsDNA [54]. More recently, binding of CTD to short 7-mer RNA and DNA oligonucleotides was also detected, although the interactions were generally weak [25]. We tested a ssDNA sequence correspondence to the ssTRS RNA used in the CTD, CTD+C-IDR, and C-IDR binding assay. Titration spectra showed no detectable binding of either the CTD or the C-IDR to ssDNA, suggesting specific binding to RNA rather than DNA (Supplementary Fig. S7A,B).

### LH negatively modulates the RNA-binding ability of CTD

Titration of LH+CTD_210-365_ with the 8-mer TRS ssRNA leads to additional peaks in the central region of the spectrum, corresponding to the disordered LH region, which are absent in the LH+CTD_210-365_ spectra (Fig. 5A). This indicates that the LH region binds RNA and undergoes substantial structural rearrangement upon RNA binding, consistent with previous reports showing that residues 219-230 of LH form an ordered helix in the presence of RNA [41]. In contrast, perturbations of CTD residues were weaker in LH+CTD_210-365_ than in isolated CTD or CTD+C-IDR (Fig. 5C) (Supplementary Fig S8A, B) . While CTD signals showed only weak perturbations, most LH peaks appeared upon RNA titration of LH+CTD at high RNA concentrations (1:5 ratio) (Fig. 5B). These results suggest that the LH negatively modulates CTD binding to RNA, whereas the C-IDR enhances it.

**Figure 5.**
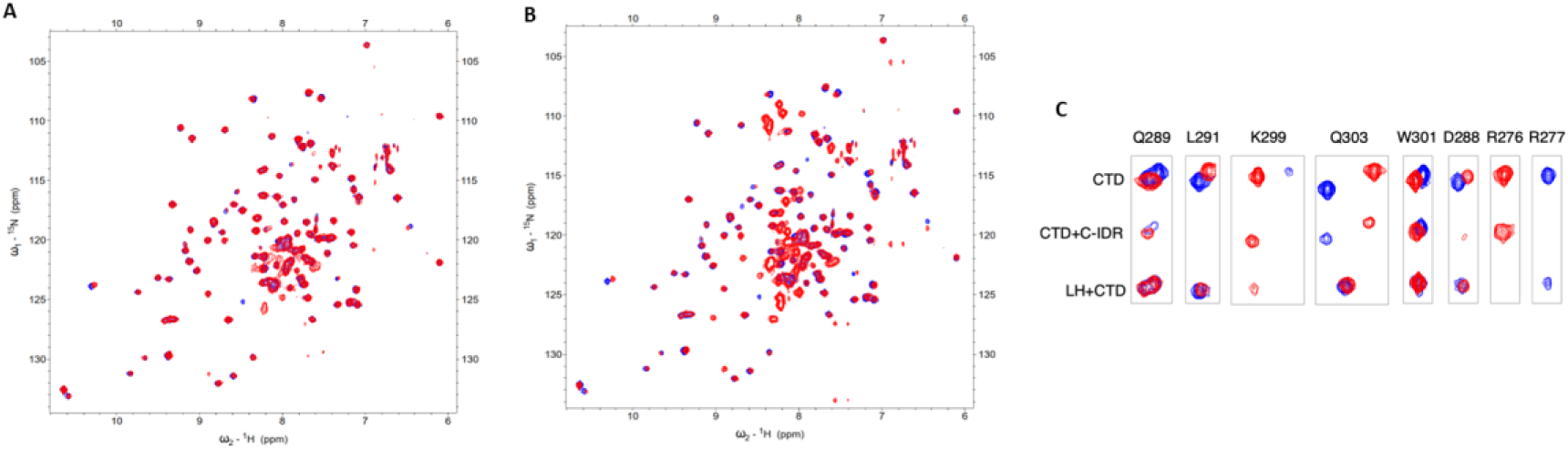
RNA binding to LH+CTD. HSQC NMR spectra of ¹⁵N-labeled constructs in the absence (blue) and presence (red) of TRS2 RNA at indicated Protein:RNA molar ratios: (**A**) LH+CTD_210-365_ with TRS RNA at 1:2 showing reappearance of LH Peaks; (**B**) LH+CTD_210-365_ with TRS RNA at 1:5, showing progressive perturbations in LH and CTD peaks with increasing TRS RNA ratio; (**C**) Comparison of the residues showing notable changes in signal intensity CTD_247-365_, CTD+C-IDR_247-419_ and LH+CTD_210-365_.

### The LLPS of CTD and C-IDR variants

Previous studies have established that LLPS of the N-protein is driven by RNA and is critical for RNA packaging, viral assembly, and infectivity [12, 57–59]. To probe the contribution of specific residues in homo-oligomerization and RNA binding within the CTD and C-IDR, we generated full-length N-protein variants carrying the mutations N_DQR_, N_1-389_, and N_R277A_. These mutant proteins were purified to homogeneity and subsequently analyzed for their phase separation behavior in the presence of an 8-mer TRS and 32-mer stem loop RNAs. For both the wild-type and mutant N-proteins, purification was carried out under high-salt conditions (1 M NaCl) with an additional heparin chromatography step to reduce potential bacterial RNA contamination that could otherwise affect LLPS (Supplementary Fig. S9). All mutant proteins, as well as wild-type N-protein, eluted as a single peak in SEC consistent with a dimeric state (Supplementary Fig. S2D) and (Supplementary Table 6). Negative-stain EM analysis further confirmed that these preparations contained only smaller-sized particles and lacked signs of RNA contamination (Supplementary Fig. S10).

In the absence of RNA, neither wild-type nor mutant N-proteins showed appreciable LLPS, highlighting the requirement of RNA for condensate formation. Upon addition of equimolar 8-mer TRS ssRNA, wild-type N-protein rapidly assembled into small, spherical condensates that gradually increased in size over 60 min (Fig. 6A).

**Figure 6.**
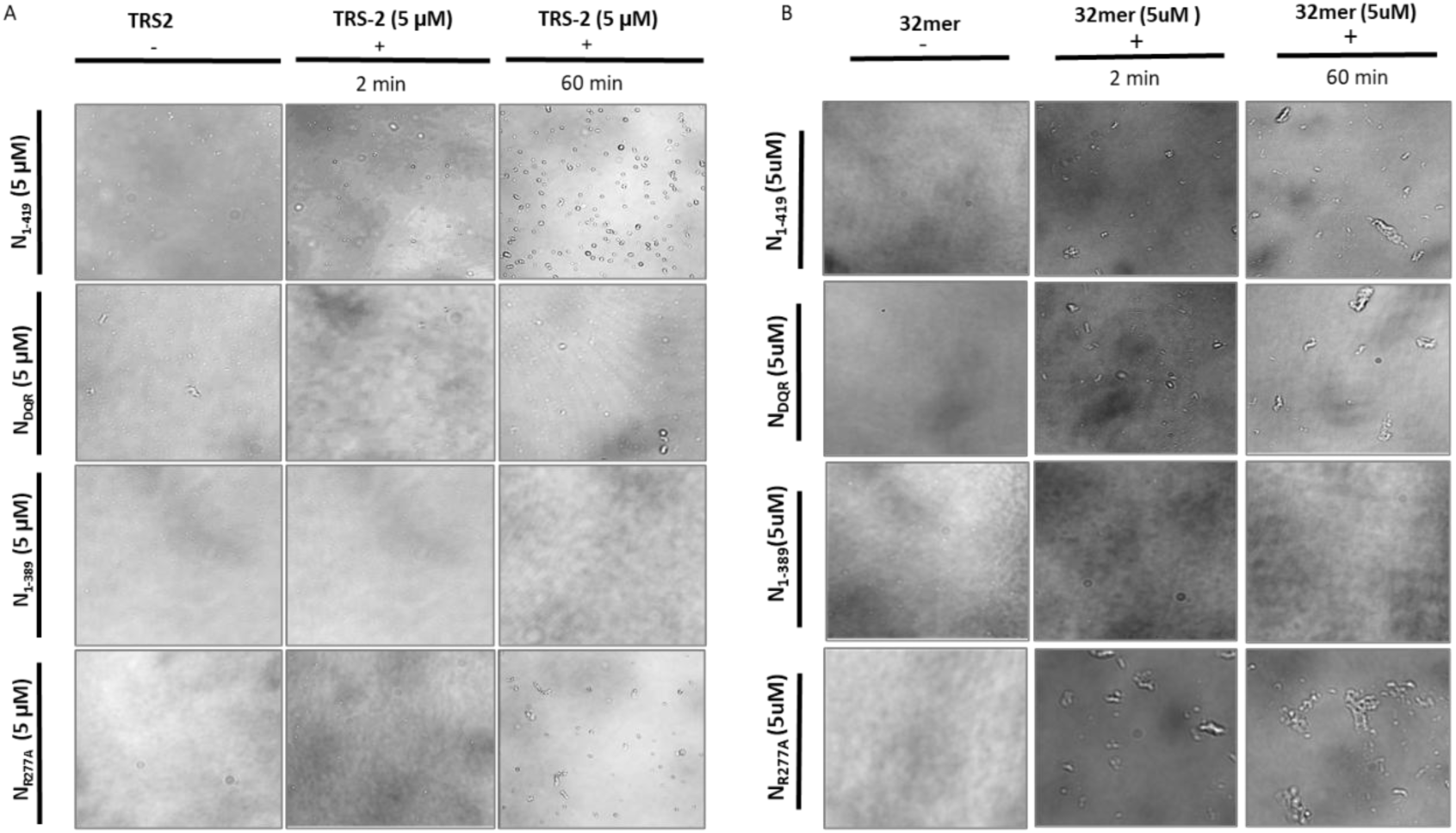
Effect of N protein mutations on LLPS (**A**) Phase seperation of N_1-419_ , N_DQR_ , N_R277A_ , and N_1-389_ in the absence(-) and presence(+) of TRS RNA at two different time points illustrating the kinetics of droplet formation; (**B**) Phase seperation of N_1-419_ , N_DQR_ , N_R277A_ , and N_1-389_ in the absence(-) and presence(+) of 32mer stem loop RNA at two different time points.

The mutant proteins showed distinct phenotypic outcomes in the presence of RNA. N_DQR_ exhibited a strongly reduced capacity to undergo LLPS, forming only small and sparse condensates compared with the wild-type N-protein. The N_1-389_ mutant was almost completely defective in RNA-induced LLPS. By contrast, the RNA-binding mutant N_R277A_ was able to form condensate; however, the droplets matured into fibril-like assemblies over time (Fig. 6A). These observations are consistent with the negative-stain EM images of the full-length N_WT and mutant proteins in the presence of RNA. Upon RNA addition, the full-length N-protein formed large, spherical particles of ∼180 nm in diameter, whereas smaller particles of ∼75 nm and ∼100 nm were observed for the N_DQR_ and N_R277A_ mutants, respectively. Notably, no spherical particles were detected for the N_1-389_ mutant (Supplementary Fig. S10). To further examine the role of longer RNA in condensate formation, we analyzed the morphology of full-length N_WT and mutant proteins in the presence of a 32-mer stem-loop RNA. Consistent with the results obtained with shorter RNAs, the 32-mer stem-loop RNA also failed to induce LLPS in the N_1–389_ mutant, indicating that deletion of the C-IDR region involved in homo-oligomerization abolishes RNA-induced condensate formation. In contrast, the addition of the 32-mer RNA led to a marked morphological transformation in the N_1-419_ , N_DQR_ , N_R277A_ constructs. Instead of forming spherical droplets, these constructs produced elongated, fibril-like condensates with longer RNA. This change in morphology suggests that RNA not only promotes phase separation but can also influence the properties and architecture of the resulting condensates, consistent with a recent report [60]. Together, these findings indicate that while RNA can trigger LLPS, multivalent protein–protein interactions are essential for condensate formation. The C-IDR homo-oligomerization interface is critical for RNA-induced LLPS, whereas the CTD oligomerization and RNA-binding interfaces primarily modulate, rather than drive, phase separation.

## DISCUSSION

Assembly of the SARS-CoV-2 nucleocapsid (N) protein with genomic RNA is driven by multivalent N–N and N–RNA interactions, yet the molecular determinants that coordinate oligomerization, RNA recognition, and LLPS remain incompletely defined. LLPS is proposed to promote RNA condensation and organize ribonucleoprotein assemblies for genome packaging [12, 27, 31, 61], and emerging evidence implicates not only the folded NTD and CTD but also the flanking IDRs in this process [37, 62, 63]. Here, we delineate the molecular basis of N-protein oligomerization with a focus on the CTD and its adjacent IDRs, the LH and C-IDR.

Our analyses reveal that while the folded CTD and individual IDR segments, LH and C-IDR, each form dimers independently, their combination markedly enhances tetramer formation, in agreement with previous observations [49]. Constructs containing both the CTD and a single IDR, such as LH+CTD or CTD+C-IDR, assemble into tetramers and exhibit a pronounced propensity for higher-order homo-oligomerization compared to the isolated LH, CTD, or C-IDR, as supported by SEC, cross-linking and NMR analyses. Inclusion of both IDRs flanking the CTD, LH+CTD+C-IDR, further enhances oligomeric assembly. Detailed characterization of the homo-oligomeric interfaces of CTD+C-IDR, and LH+CTD using NMR and mutational studies identified two oligomerization interfaces: residues centered on the CTD α2-helix, including residues D288, Q289, and R293 and residues within the C-IDR spanning positions 390-409. The functional significance of these interfaces was evaluated through EM and LLPS using full-length N protein mutants, demonstrating that disruption of either interface impairs oligomerization and RNA-dependent LLPS. In particular, deletion of the C-IDR region (N_1-389_) abolishes condensate formation, and the N_DQR_ variant diminishes LLPS, highlighting the essential contribution of the C-IDR.

These findings are consistent with lysine-crosslinking mass spectrometry, which identified both LH and C-IDR as key interaction hubs that drive N protein phase separation in the presence [40] and absence of RNA [37]. Structural studies further reveal transient helices spanning residues 219-230 in the LH [41, 55] and residues 383–396 and 402–415 in the C-IDR [55]. The LH can form trimeric or tetrameric coiled-coil–like structures that promote homo-oligomerization [62], and our results suggest that transient helices within the C-IDR play an analogous role in facilitating homo-oligomerization. Notably, mutations in SARS-CoV-2 N protein frequently accumulate within all three IDRs, the N-IDR, LKR, and C-IDR [17, 64, 65] and the mutationally protected sequence “islands” map to transient helical segments of the C-IDR residues 390–394 and 403–408 as well as in the LH residues 218–231 [64, 65]. The common variant, such as G215C, occurs near the LH enhances both homo-oligomerization and viral infectivity [65, 66]. While within the C-IDR, mutations such as D377Y, which are enriched in Delta variants, increase RNA binding and infection [67]. In contrast, acetylation or acetylation-mimicking substitutions at Lys375 disrupt N protein LLPS with RNA, emphasizing the critical role of the C-IDR [68, 69]. Other frequently observed C-IDR mutations, including T362I, A376T, E378V, D402Y, and S413R, warrant investigation to determine their effect on oligomerization, RNA binding, and LLPS [70–73]. Together, these observations support a model in which the LH, CTD, and C-IDR act as cooperative oligomerization modules that enable higher-order assembly and RNA-dependent condensate formation and suggest that mutations in these regions may modulate viral assembly, infectivity, and pathogenesis.

In addition to its primary RNA-binding NTD, the N protein harbors multiple auxiliary RNA-binding regions [25–27] . Although the CTD and its adjacent IDRs are known to interact with RNA, their precise roles in LLPS remain incomplete [74] . Our NMR titration and mutational analyses revealed a distinct RNA-binding site within the CTD, separate from the highly positively charged dimerization interface, and identified the conserved residue R277 as a key determinant of CTD–RNA recognition. Notably, the presence of the C-IDR enhanced CTD RNA binding, consistent with cooperative CTD and C-IDR interactions with RNA [25]. Correspondingly, the R277A mutation reduced RNA binding of both CTD and CTD+C-IDR, supporting the cooperative nature of their RNA binding. Although the LH region is capable of RNA binding, it attenuates CTD–RNA interactions in the LH+CTD context, supporting a regulatory role for IDRs in tuning RNA-binding specificity. Introducing the R277A mutation into the FL-N (N_R277A_) altered condensate formation without abolishing LLPS, indicating that CTD-mediated RNA binding shapes condensate architecture rather than initiating phase separation. Furthermore, the appearance of several LH and C-IDR resonances upon RNA addition suggests that RNA stabilizes transient helical elements that contribute to oligomerization and condensate formation.

In summary, our results define discrete N–N and N–RNA interaction interfaces that collectively drive higher-order assembly and RNA-induced LLPS, processes central to genome packaging and virion formation. The cooperative roles of CTD, LH, and C-IDR provide a structural framework for understanding N-protein assembly and identify potential molecular surfaces that could be exploited to disrupt coronavirus ribonucleoprotein formation.

## Acknowledgement

We thank Dr. Sreekanth Rajan for his critical suggestions on the manuscript. We thank Chandrani Samadder from Cryo-EM facility for helping us acquire TEM images. The NMR data were acquired at the National Center for Biological Science-Tata Institute of Fundamental Research NMR Facility. The confocal microscopy experiments were performed at the Central Imaging and Flow-cytometry Facility (CIFF) at the National Center for Biological Science-Tata Institute of Fundamental Research. The TEM images were acquired at the Cryo-EM facility at Institute for Stem Cell Science and Regenerative Medicine.

## Author contributions

Sneha G Bairy (Data curation [equal], Formal analysis [equal], Investigation [lead], Methodology [equal], Writing—original draft [equal], Writing—review & editing [equal]), Thazhe Kootteri Prasad (Data curation [equal], Formal analysis [equal], Investigation [lead], Methodology [equal], Writing—original draft [equal], Writing—review & editing [equal]), Yogeshwar Saravana Kumar (Data curation [equal], Formal analysis [equal], Investigation [lead], Methodology [equal], Writing—review & editing [equal])), B Ganavi (Investigation [supporting], Methodology [supporting]), Shwetha S (Investigating [supporting], Methodology [supporting]), Saranya S (Investigation [supporting], Methodology [supporting]), Sriram Prakash B (Investigation [supporting]) Vignesh Sounderrajan (Investigation [supporting], Methodology [supporting]), Krupakar Parthasarathy (Funding acquisition [lead], Investigation [supporting], Methodology [supporting], Writing—review & editing [lead]). Neelagandan Kamariah (Conceptualization [lead], Formal analysis [equal], Funding acquisition [lead], Investigation [supporting], Methodology [supporting], Project administration [lead], Supervision [lead], Writing—original draft [lead], Writing—review & editing [lead]).

## Supplementary data

Supplementary data is available NAR online

## Data availability

The data that support the findings of this study are available from the corresponding author upon reasonable request.

