## SUPPLEMENTARY INFORMATION for "Molecular Characterization of SARS-CoV-2 N Protein Interfaces: Implications for Oligomerization, RNA Binding, and Phase Separation"

### SUPPLEMENTARY TABLES

**Supplementary Table 1**. List of primers used in this study

| | |
|---|---|
| 362_NPC_247_F | GAA GGA GAT ATA CC ACG AAG AAA AGC GCG GCT G |
| 363_NPC_365_F | GAA GGA GAT ATA CCccaacagagcctaagaaggac |
| 364_NPC_vec_r (pet28_r_without N his) | GGTATATCTCCTTCTTAAAGTTAAACAAA |
| 368_NPC_247_F | GAA GGA GAT ATA CC ACG AAG AAA AGC GCG GCT G |
| 369_NPC_365_F | GAA GGA GAT ATA CCccaacagagcctaagaaggac |
| 370_NPC_vec_r | GGTATATCTCCTTCTTAAAGTTAAACAAA |
| 371_NPC_247_f2 | GAA GGA GAT ATA CC atg ACG AAG AAA AGC GCG GCT G |
| 372_NPC_365_f2 | GAA GGA GAT ATA CC atg ccaacagagcctaagaaggac |
| p382_N247_pet_f | TACTTCCAGGGaTCCagc atgacgaagaaaagcgcggctg |
| p383_N365_pet_r | ctcgacttaCTCGAGttaaggaaaagtcttgtaggcatcgatG |
| p425_Nucleocapsid_399_r | gtgctcgacttaCTCGAGttaatccgcggcgggaaggag |
| p442_Nucleocapsid_389_r | gtgctcgacttaCTCGAGttattgcttcttttgacgttggggtag |
| p443_Nucleocapsid_379_r | gtgctcgacttaCTCGAGttaagtttcgtcagctttcttcttcttgtc |
| p444_Nucleocapsid_369_r | gtgctcgacttaCTCGAGttacttaggctctgttggaggaaaag |
| p423_Nucleocapsid_409_r | gtgctcgacttaCTCGAGttactgttgcagctgtttcgagaaatc |
| p425_Nucleocapsid_399_r | gtgctcgacttaCTCGAGttaatccgcggcgggaaggag |
| p455_Nucleocapsid_374_r | gtgctcgacttaCTCGAGttacttcttcttgtccttcttaggctc |
| P494_N_D288_A_f | cgga gcc caagagttaattaggcagggtacg |
| P495_N_288_A_r | ctctcttgggctccgaaattaccctgcgtttgttccgG |
| P496_N_R293_A_f | G TTA ATT gcg cagggtacggactacaagcattg |
| P497_N_R293_A_r | ccctgcgcAATTAACTCTTGGGCTCCGAAATTACC |
| P498_N_Q_289_A_f | GGA GCC GCA GAG TTA ATT AGG CAG GGT ACG GAC |
| P499_N_Q_289_A_r | TTAACTCTGCGGCTCCGAAATTACCCTGCGTT TG |
| P505_N_R277A_f | GGAAGAGCCGGTCCGGAACAAACGCAGG |
| P506_N_R2771_r | GGACCGGCTCTTCCAAAAGCCTGTGTAACATT GTAC |
| P533_N_R293A_f | GT TAA TT GCA CAG GGT ACG GAC TAC AAG CATTG |
| P534_N_R293A_r | CCCTGtgcAATTAACTCTGCGGCTCCGAAATTA C |
| p560_N_210_f | GAATTTATACTTCCAGGGaTCC atggcaggcaatggaggggatg |

**Supplementary Table 2**. List of constructs made in this study

| **N_Protein constructs** | **Amino acid residue boundaries/ Mutations** |
|---|---|
| $N_{1-419}$ | Residues 1-419 |
| $N_{DQR}$ | Residues 1-419 with mutations D288A, Q289A, R293A residues |
| $N_{1-389}$ | Residues 1-389 |
| $N_{R277A}$ | Residues 1-419 with R277A mutation |
| $CTD_{247-365}$ | Residues 247-365 |
| $CTD+C\text{-}IDR_{247-419}$ | Residues 247-419 |
| $CTD+C\text{-}IDR_{247-409}$ | Residues 247-409 |
| $CTD+C\text{-}IDR_{247-399}$ | Residues 247-399 |
| $CTD+C\text{-}IDR_{247-389}$ | Residues 247-389 |
| $C\text{-}IDR_{366-419}$ | Residues 366-419 |
| $CTD_{R277A}$ | Residues 247-365 with R277A mutation |
| $CTD+C\text{-}IDR_{R277A}$ | Residues 247-419 with R277A mutation |
| $CTD+C\text{-}IDR_{DQR}$ | Residues 247-419 with D288A, Q289A, R293A mutations |
| $LH+CTD_{210-265}$ | Residues 210-365 |
| $LH+CTD_{210-419}$ | Residues 210-419 |

**Supplementary Table 3.** Estimation of molecular weights and oligomeric states of N protein constructs

| N_Protein constructs | Calculated MW from sequence (kDa) | Apparent MW from SEC (kDa) | Estimated Oligomeric state |
|---|---|---|---|
| $CTD_{247\text{-}365}$ | 13 | 36 | 2.7 |
| $LH+CTD_{210\text{-}365}$ | 17 | 64 | 3.7 |
| $CTD+C\text{-}IDR_{247\text{-}419}$ | 19 | 87 | 4.5 |
| $LH+CTD+C\text{-}IDR_{210\text{-}419}$ | 23 | 96 | 4.1 |
| $C\text{-}IDR_{366\text{-}419}$ | 6 | 19 | 3.0 |
| $N_{1\text{-}419}$ | 48 | 141 | 2.9 |

**Supplementary Table 4.** Estimation of molecular weights and oligomeric states of CTD+C-IDR mutants

| N Protein constructs | Calculated MW from sequence (kDa) | Apparent MW from SEC (kDa) | Estimated Oligomeric state |
|---|---|---|---|
| $CTD+C\text{-}IDR_{247–419}$ | 19 | 87 | 4.5 |
| $CTD+C\text{-}IDR_{DQR}$ | 19 | 76 | 4.0 |
| $CTD+C\text{-}IDR_{247–409}$ | 18 | 86 | 4.6 |
| $CTD+C\text{-}IDR_{247–399}$ | 17 | 66 | 3.8 |
| $CTD+C\text{-}IDR_{247–389}$ | 16 | 63 | 3.9 |
| $CTD+C\text{-}IDR_{247–379}$ | 15 | 53 | 3.5 |

**Supplementary Table 5.** The percentage helical content in $C\text{-}IDR_{366\text{-}419}$ was estimated using BeStSel

| Concentration | Helix % |
|---|---|
| 50uM | 0 |
| 100uM | 5 |
| 200uM | 12 |
| 400uM | 15 |

**Supplementary Table 6.** Estimation of molecular weights and oligomeric states of FL-N and its mutants

| N_Protein constructs | Calculated MW from sequence (kDa) | Apparent MW from SEC (kDa) | Estimated Oligomeric state |
|---|---|---|---|
| 1. $N_{1\text{-}419}$ | 48 | 141 | 2.9 |
| 2. $N_{R277A}$ | 48 | 101 | 2.0 |
| 3. $N_{DQR}$ | 48 | 128 | 2.6 |
| 4. $N_{1\text{-}389}$ | 45 | 153 | 3.4 |

### SUPPLEMENTARY FIGURES

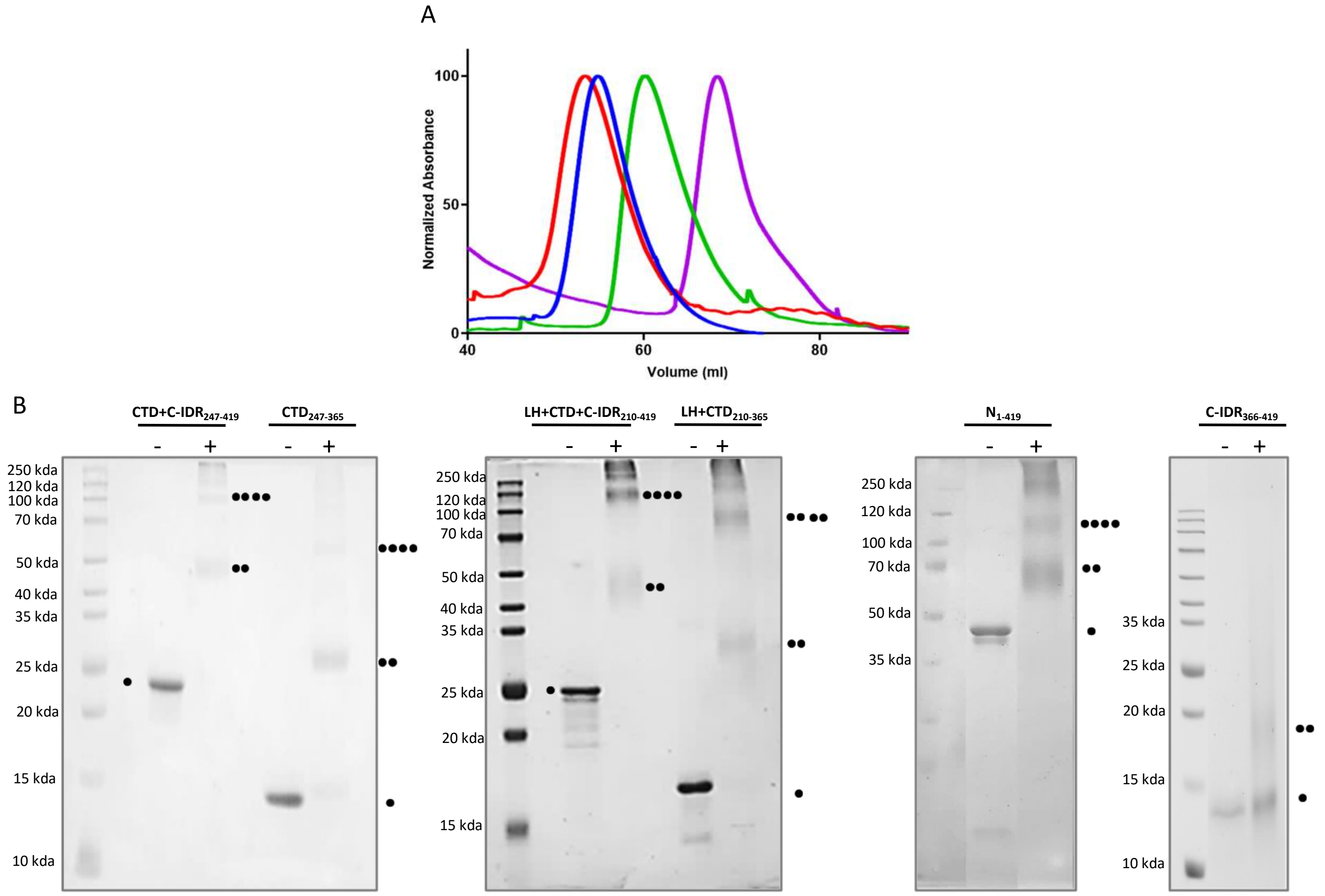


**FigureS1**. Oligomeric state of CTD flancked by IDR's (**A**) Comparison of SEC elution profiles for LH+CTD+C-$IDR_{210\text{-}419}$ (red), LH+$CTD_{210\text{-}365}$ (green), CTD+C-$IDR_{247\text{-}419}$ (blue), and $CTD_{247\text{-}365}$ (purple), highlighting differences in elution volume and oligomeric state across the constructs; (**B**) Glutaraldehyde cross-linking analysis of CTD+C-IDR247−419, $CTD_{247\text{-}365}$, LH+CTD+C-IDR210−419, LH+CTD210−365 and C-IDR366-419 in the absence (-) and presence (+) of 0.3% glutaraldehyde, illustrating their oligomeric state. Symbol denote Monomer (●), dimer (●●), tetramer (●●●●).

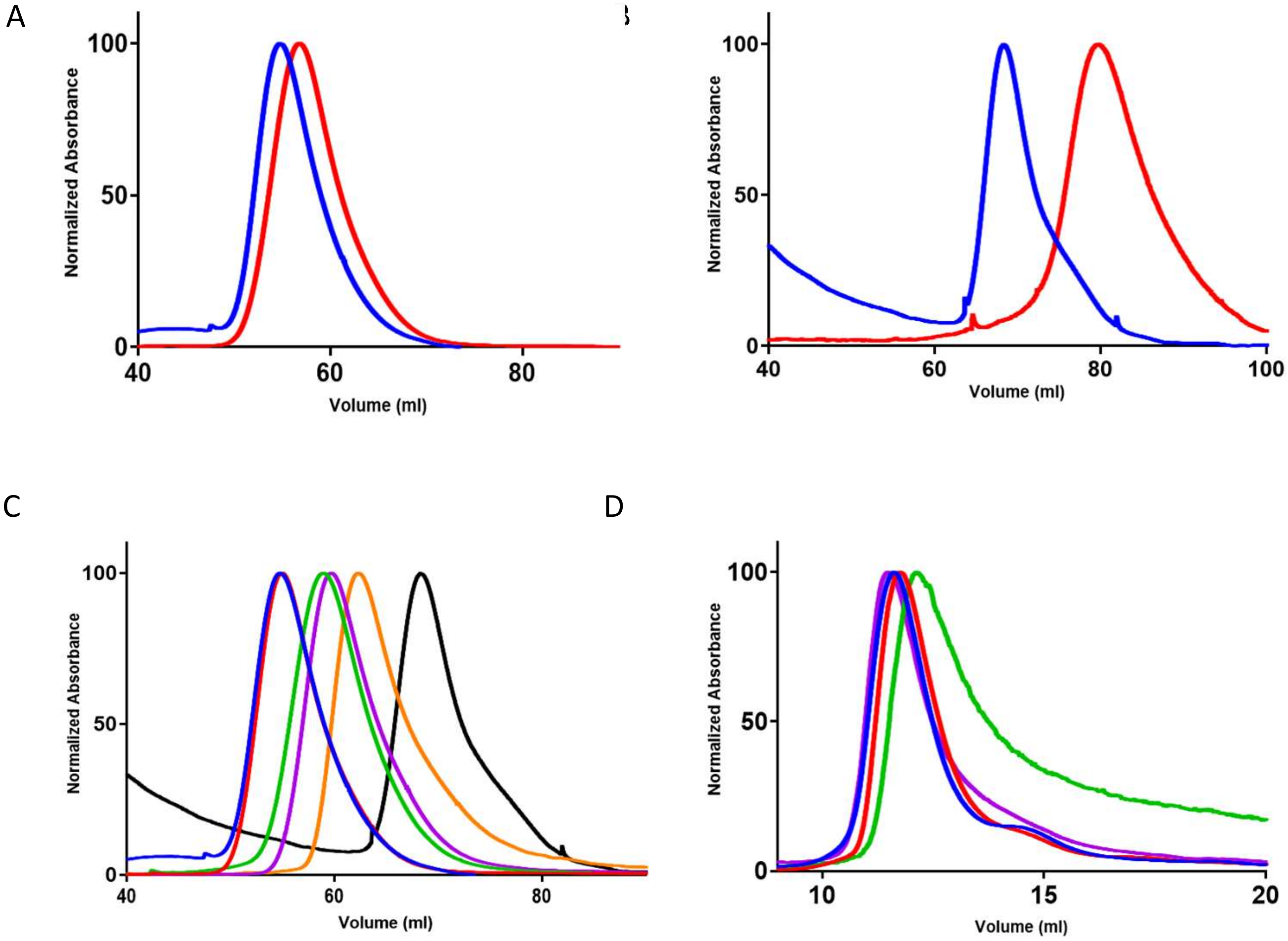


**Figure S2.** SEC elution profiles of various N constructs (**A**) Comparison of SEC profiles for CTD+C-IDR$_{247\text{-}419}$ (blue), and CTD+C-IDR$_{DQR}$ (red), (**B**) Comparison of SEC profiles for CTD$_{247\text{-}365}$ (blue), and IDR$_{366\text{-}419}$ (red), (**C**) Comparison of SEC profiles for CTD+C-IDR$_{247\text{-}419}$ (blue), CTD+C-IDR$_{247\text{-}409}$ (red), CTD+C-IDR$_{247\text{-}399}$ (green), CTD+C-IDR$_{247\text{-}389}$ (purple), CTD+C-IDR$_{247\text{-}379}$ (orange), and CTD$_{247\text{-}365}$ (black) and (**D**) Comparison of SEC profiles for N$_{1\text{-}419}$ (blue), N$_{DQR}$ (red), N$_{R277A}$ (green), and N$_{1\text{-}389}$ (purple).

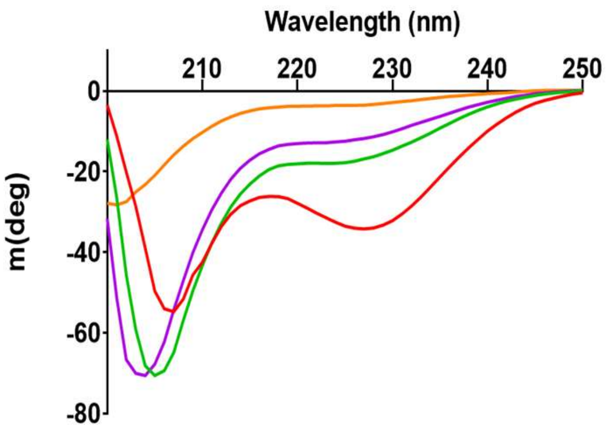


**Figure S3.** Concentration-dependent helix formation of C-IDR$_{366\text{-}419}$ : The CD spectra of C-IDR showed an increase in helicity in a concentration-dependent manner. Spectra were recorded at at 50 μM (orange), 100 μM (purple), 200 μM (green) and 400 μM (red).

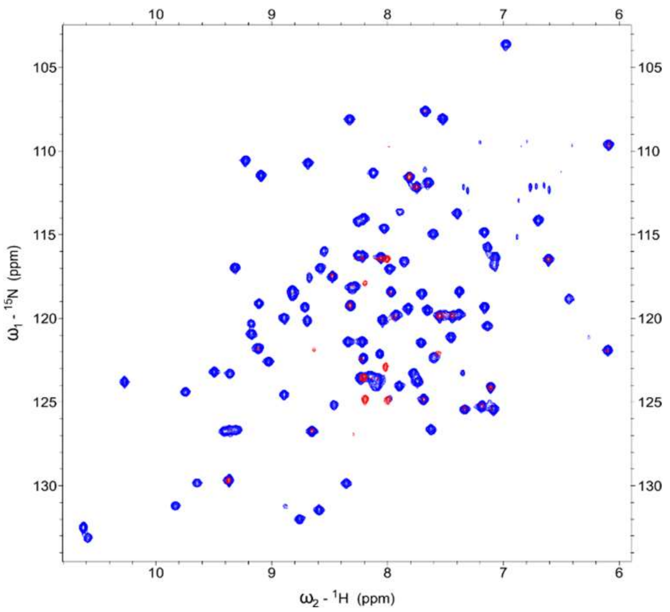


**Figure S4**. HSQC NMR spectra of $^{15}$N-labeled CTD$_{247\text{-}365}$ in the absence ( blue), in the presence (red) of 32-mer RNA at a 1:0.05 molar ratio.

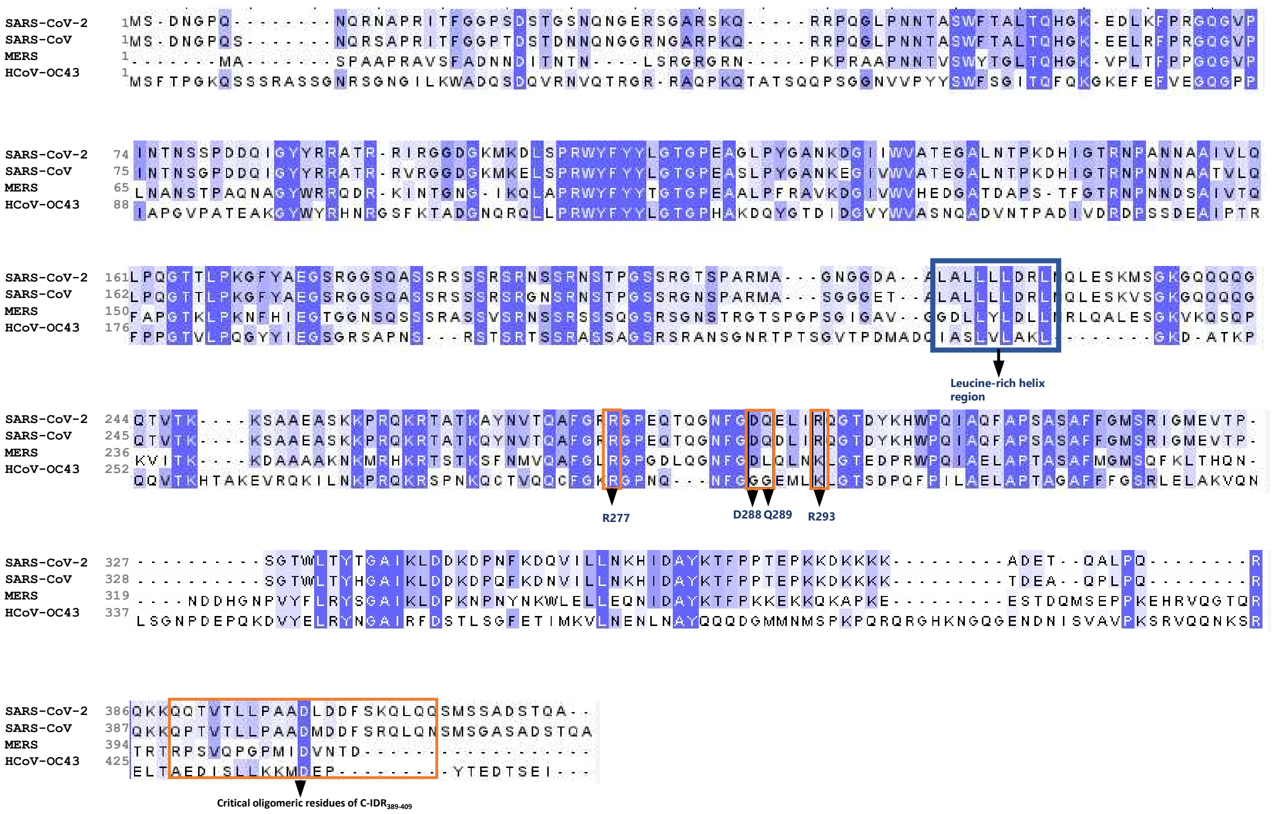


**Figure S5**. Sequence comparison of betacoronavirus N proteins (A) Multiple sequence alignment showing that residue R277 within the CTD is highly conserved.

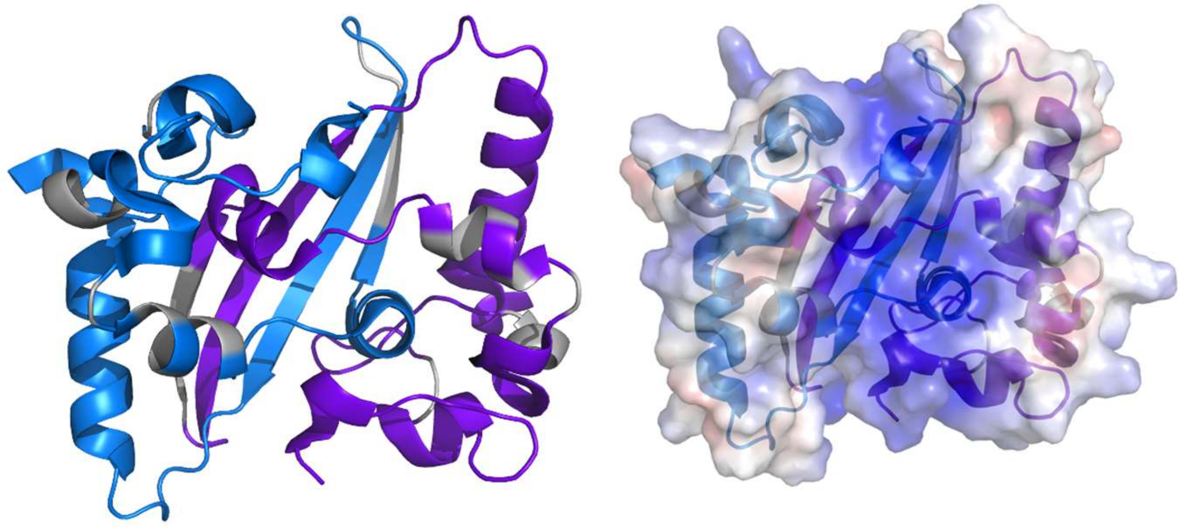

**Figure S6**. RNA-binding interface of the CTD. The dimeric structure of the CTD is shown with each monomer depicted in a different color, and the TRS ssRNA-binding interface highlighted in grey. The electrostatic surface representation of the SARS-CoV-2 CTD dimer reveals a prominently positively charged region on the helical face of the rectangular slab-like dimer.

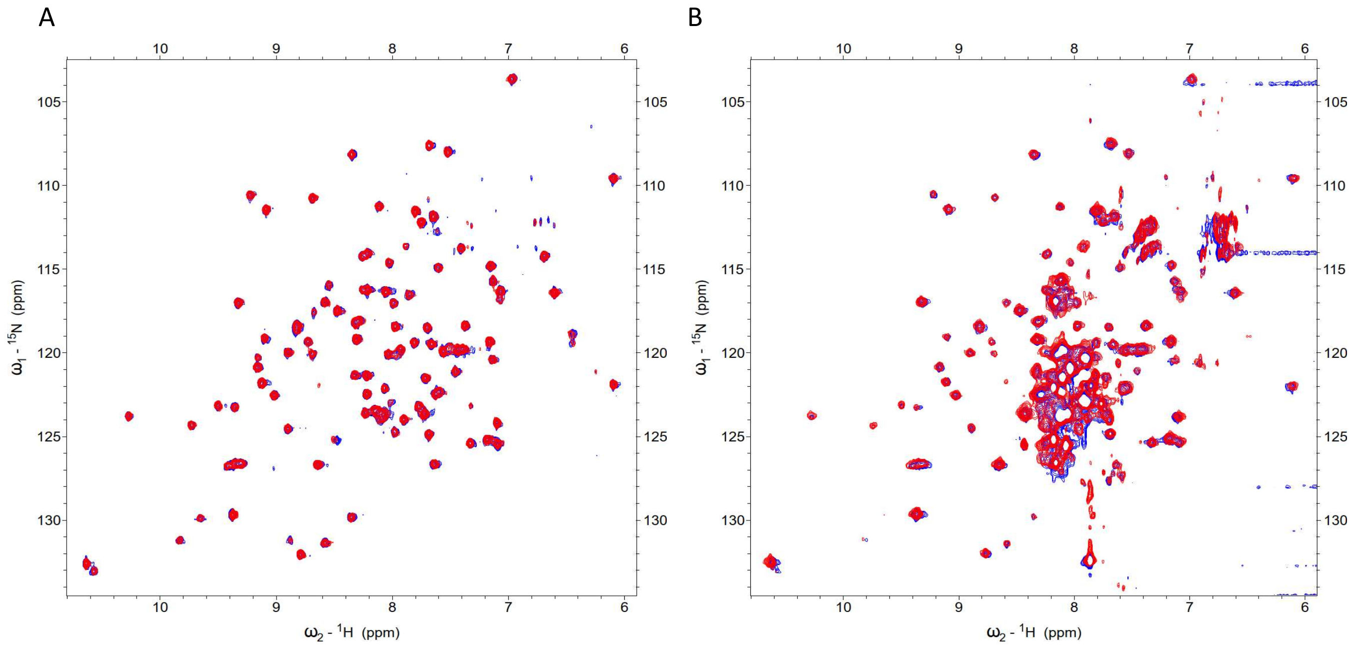


**Figure S7**. HSQC NMR spectra of $^{15}N$-labeled constructs in the absence (blue) and presence (red) of DNA corresponding to the TRS RNA sequence at the indicated Protein: DNA molar ratio. (**A**) $CTD_{247-365}$ with DNA (1:5); (**B**) CTD+C-$IDR_{247-419}$ with DNA at 1:1 molar ratio, showing no significant interaction.

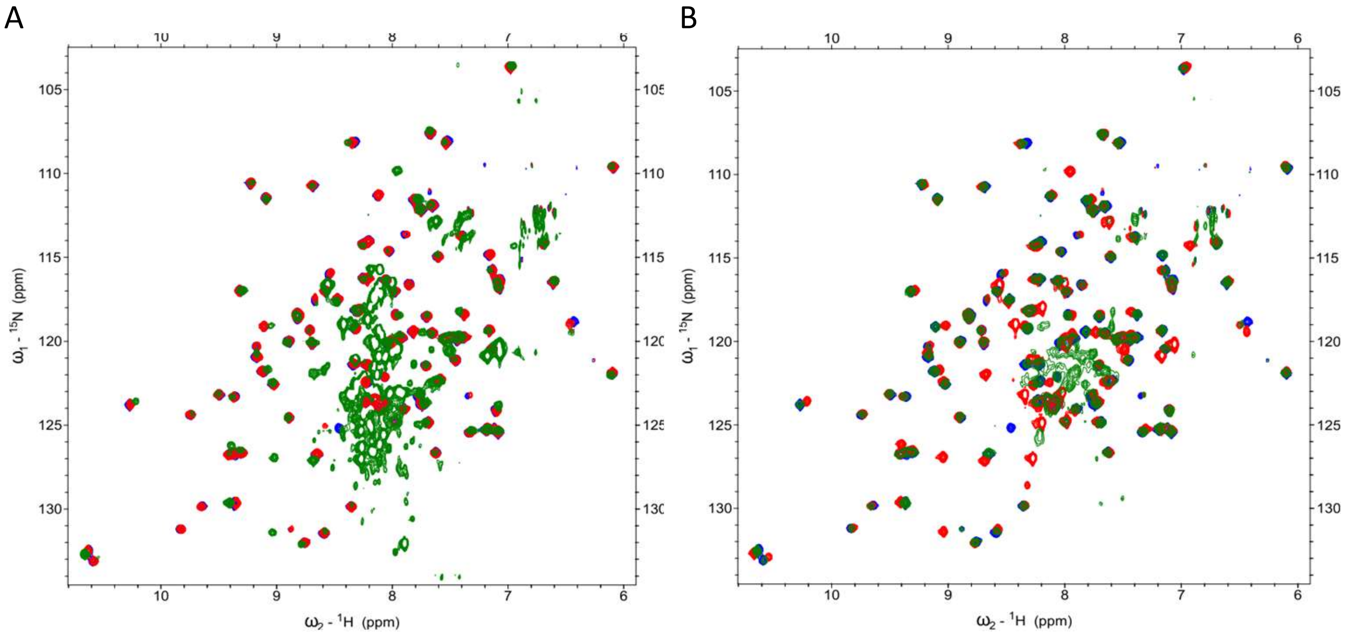


**Figure S8**. HSQC NMR spectra of $^{15}$N-labeled constructs in the absence ($CTD_{247\text{-}365}$, blue), (**A**) in presence of TRS RNA (1:1) ($CTD_{247\text{-}365}$ ,red), and ($CTD\text{+}C\text{-}IDR_{247\text{-}419}$ ,green); (**B**) in presence of TRS RNA (1:2) ($CTD_{247\text{-}365}$ ,red), and ($LH\text{+}CTD_{210\text{-}365}$ ,green).

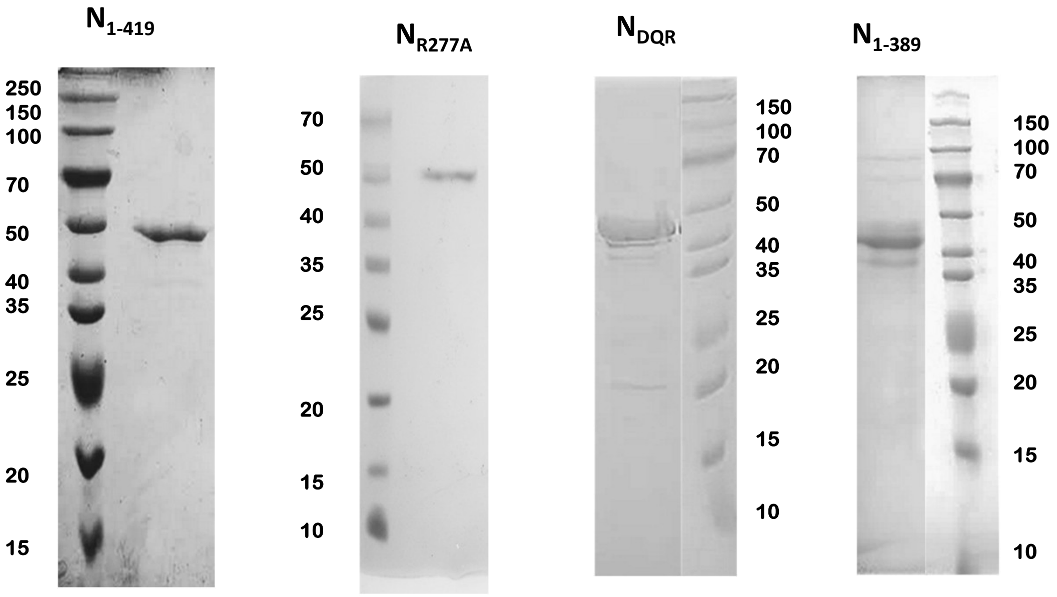


**Figure S9**. SDS-PAGE analysis of purified full-length N protein and its mutants

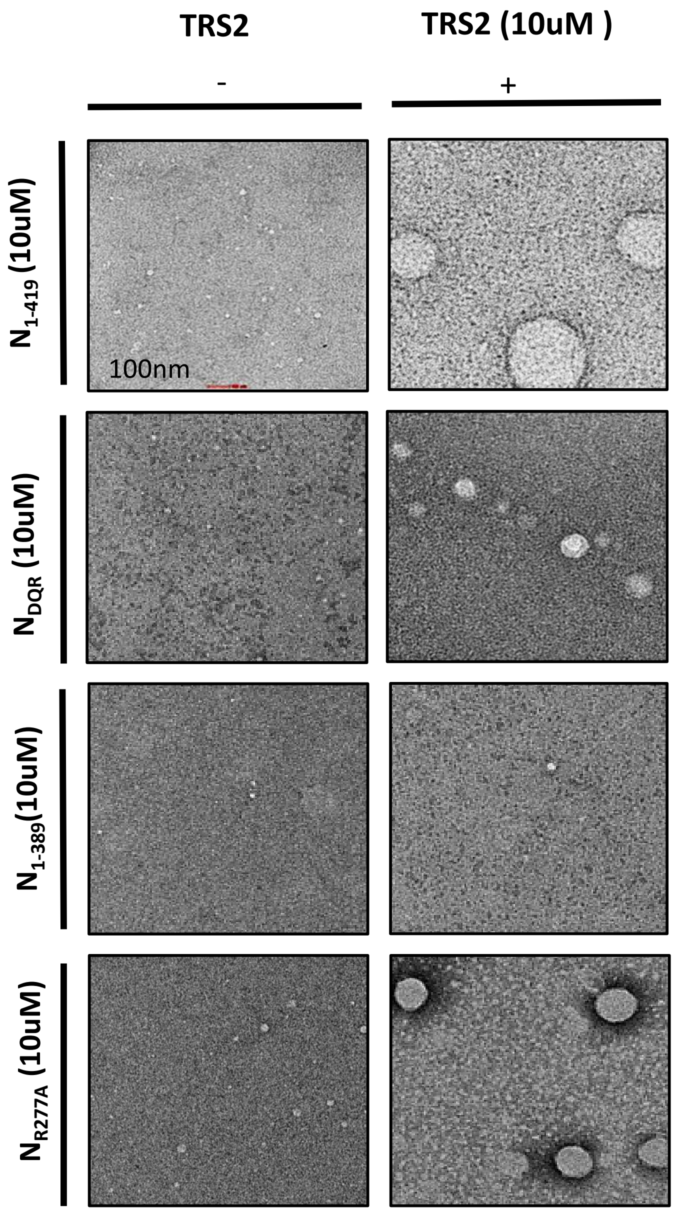


**Figure S10**. Negative staining of $N_{1\text{-}419}$ , $N_{DQR}$ , $N_{R277A}$ , and $N_{1\text{-}389}$ in the absence(-) and presence of (+) TRS RNA
